# Effects of Ambient Temperature on the Recognition and Preferences for Host Song Pulse-rate by the Acoustic Parasitoid Fly *Ormia ochracea*

**DOI:** 10.1101/2023.03.29.534770

**Authors:** Karina J. Jirik, Jimena A. Dominguez, Iya Abdulkarim, Johanna Glaaser, Emilia Stoian, Luis J. Almanza, Norman Lee

## Abstract

Receivers of acoustic communication signals evaluate signal features to correctly identify conspecifics. Environmental variation such as changes in the ambient temperature can alter signal features that may render species recognition a challenge. In field crickets, song temporal features evaluated for species recognition vary with temperature, and the intended receivers of these signals exhibit signal preferences that are temperature-coupled to maintain effective communication. Whether eavesdroppers of communication signals exhibit similar temperature-coupled preferences is unknown. Here, we investigate whether the parasitoid fly *Ormia ochracea*, an eavesdropper of cricket calling songs, exhibit song pulse rate preferences that are temperature-coupled. We use a high-speed treadmill system to record walking phonotaxis at three ambient temperatures (21, 25, and 30 °C) in response to songs that varied in pulse rates (20 to 90 pulses per second). Total walking distance, peak steering velocity, angular heading, and the phonotaxis performance index varied with song pulse rates and were affected by ambient temperature. The peak of phonotaxis performance index preference functions became broader and exhibited a “high-pass” shape, shifting to higher pulse rate values at higher temperatures. Temperature related changes in cricket songs between 21-30 °C will not drastically affect the ability of flies to recognize cricket calling songs.

## Introduction

A complete understanding of the evolution of communication systems requires in-depth knowledge of how environmental factors influence the behavior of signalers and receivers [1]. Ambient temperature is one such environmental factor that can vary in time and space, with profound effects on locomotion, reproduction, foraging, and communication behavior of animals [2,3]. This is especially the case for ectotherms where body temperature is regulated by the temperature of the environment. Behaviors that are modulated by temperature can occur over a continuum that span short to long term effects. In the short term, higher ambient temperature may elevate metabolic rates [4] and can lead to increased foraging behavior [5] to meet increased metabolic demands. Deviations from an optimal temperature range can reduce locomotor performance and limit the ability of animals to locate food [6–8]. Similar temperature-dependent effects have also been found to alter the production of temporally patterned communication signals [9–12], the transmission of signals through the environment [13], or the functioning of sensory systems [14–16], all of which can impact reproductive success [17,18].

Production of acoustic communication signals involve rhythmic motor activity. Changes in muscle physiology resulting from changes in ambient temperature can affect the temporal [19–29] and spectral [17,26,28,30] features of advertisement calls in a range of animals. In field crickets, males communicate with acoustic signals that are produced by stridulation, a process that involves rubbing together specialized structures on the forewings. Contraction of forewing muscles opens and closes the left and right forewings [31,32]. Each wingstroke closure generates a sound pulse that is created by the plectrum of the cricket’s left forewing striking subsequent pegs that form the file of the cricket’s right forewing. Wingstroke closures occur at what is typically a species-specific song pulse rate and this rate is multiplied by the scraper engaging subsequent teeth in the file. This excitation is transduced to the harp of the forewings where it resonates at the carrier frequency of the song [33]. As song pulse rate is determined by the rate of muscle contractions, pulse rates can vary depending on ambient temperature [20,21,34].

Temperature coupling occurs when temperature-dependent changes in signal characteristics are matched with parallel changes in signal preferences exhibited by intended receivers [22,25,29,30,35]. This temperature coupling ensures that signal recognition is maintained, and that receivers of courtship signals can maintain selectivity for high quality mates, which can allow for sexual selection to operate across a range of ambient temperatures. Accordingly, temperature coupling may be an adaptive function to ensure that thermal changes in signal production are matched with receiver signal recognition and preferences. In the field cricket *Gryllus firmus*, the chirp rate and pulse rate of calling songs vary with temperature, and females exhibit a parallel change in preferences when these features vary with temperature [25]. One proposed explanation for temperature coupling is that signalers and receivers may share common neural elements due to genetic coupling [36]. Receivers may possess silent neural circuitry that is homologous to central pattern generators for song production in males. Auditory input would elicit corollary discharges in the silent central pattern generator, which may be compared to a male song template at higher levels of the auditory system. While this might be a plausible explanation for intended receivers that are genetically coupled to the signaler [37], whether and how unintended receivers might exhibit signal preferences that parallel temperature dependent changes in song features remains to be examined.

The acoustic parasitoid fly *Ormia ochracea*, an unintended receiver of cricket calling songs, rely on directional hearing to eavesdrop on the calling songs in search of appropriate host species [38–40]. With ears that are separated by no more than 500μm [41], the small physical size of *O. ochracea* severely limits directional cues that are available from a 5 kHz calling song [42]. Directional hearing in *O. ochracea* is made possible with a pair of mechanically coupled eardrums that function to amplify almost non-existing directional cues in the sound field, to magnitudes that are amenable to neuronal processing [43,44]. This specialized mechanical apparatus is coupled with an auditory nervous system that exhibits exquisite sensitivity to temporal information [45]. Most of the primary auditory afferents are type 1 receptors, which exhibit phasic, low jitter spikes that are timed with the onset of sound pulses and are ideal for copying the temporal pattern of cricket songs [45]. Song recognition is based on evaluating the temporal patterning of sound pulses [40,46]. In Florida, *Gryllus rubens* is the preferred cricket host [47,48], which produces calling songs characterized by a pulse rate of about 45-50 pulses per second (pps). Floridian *O. ochracea* exhibit pulse rate preference functions that peak at the same range of pulse rates [49]. Upon detection of appropriate calling songs, gravid female flies will perform flying and walking phonotaxis to the source location [50–52], where they deposit first instar planidia near or on top of the host. The planidia will then burrow into the cricket, develop for a period of 7 to 10 days, emerge from the cricket, pupate, and finally hatch as an adult fly [53–55]. Reproduction in *O. ochracea* crucially depends on the ability of flies to recognize and localize host cricket calling songs.

Whether temperature-dependent changes in song pulse rates affect the ability of *O. ochracea* to recognize and localize appropriate host cricket calling songs is currently unknown. While *O. ochracea* are not genetically coupled to crickets, gravid female flies are expected to evolve signal preferences that parallel thermal-induced changes in signal features. In this study we specifically test the hypothesis that preferences for signal features exhibited by eavesdroppers are coupled to temperature-dependent changes in signal features. According to this hypothesis, we expect that as the pulse rate of calling songs increases with ambient temperature, the peak in pulse rate preference functions exhibited by Floridian *O. ochracea* will show a concomitant shift to higher pulse rate values. To address this hypothesis, we measured walking phonotactic responses of *O. ochracea* when subjected to different ambient temperatures in response to calling songs that varied in pulse rates. Our results indicate that the peak of pulse rate preference functions in Floridian *O. ochracea* shift with ambient temperature. Pulse rate preference functions also appear to broaden at higher temperatures, indicating lower levels of pulse rate selectivity at higher temperatures.

## Materials and Methods

### Animals

Behavioral walking phonotaxis experiments were conducted on gravid female *Ormia ochracea* from a laboratory colony that was originally derived from Gainesville, Florida. Flies were reared in a temperature, humidity, and light controlled environmental chamber (Power Scientific, Inc., Model DROS52503, Pipersville, PA) set to a 12h:12h light/dark cycle, 25 °C, at 75% humidity, and provided butterfly nectar solution (The Birding Company, MA) *ad libitum*.

### Experimental Setup

Walking phonotactic responses were recorded from flies tethered and held on top of a spherical treadmill system previously described in [56] and [49]. This treadmill system consists of a spherical ping-pong ball held afloat in a custom holder with a continuous air stream. Rotations of the spherical treadmill actuates a modified optical mouse sensor (ADNS 2620, Avago Technologies, USA) that is situated directly below the treadmill. This optical mouse sensor registers motion as changes in x and y pixel units at a sampling rate of 2160 Hz [56]. In a previous study [49], we confirmed that *O. ochracea* never caused the spherical treadmill to rotate on its axis and thus a single sensor system is sufficient for recording *O. ochracea* walking phonotactic responses. Data captured from the treadmill system was synchronized with sound presentation using software (StimProg V6) developed in MATLAB (R2018a, The MathWorks Inc., USA) that interfaced with National Instruments Data Acquisition hardware (NI USB-6363, USA). Changes in x and y pixel values were calibrated to real-world distances in centimeters by measuring the actual displacement of the spherical treadmill in footage captured using a high-speed camera (Chronos 1.3 High-speed Camera, 1,000 frames per second, Krontech, Canada). Images from this high-speed camera were imported into ImageJ (ver 1.51), and the displacement of markers on the spherical treadmill were measured relative to a known distance. This treadmill system was situated 25 cm away from two silk-dome tweeters (1–1/8 Dayton Audio Classic Series DC28FS-8, USA), which were located at ±45° azimuth (to the left and right of the treadmill system) relative to the forward (0° azimuth) direction of a tethered fly. This experimental setup was situated on a vibration isolation table (TMC) and located in an acoustically dampened sound chamber (Wenger Soundlok, USA). Ambient temperature in the sound chamber was maintained using a combination of a Lasko space heater (model CT2272, China) and the building HVAC system, and was verified with a thermometer located directly above the treadmill system.

### Acoustic stimuli

Acoustic stimuli consisting of synthetically produced cricket songs that were created in MATLAB. These cricket songs were composed of tonal sound pulses with a carrier frequency of 5 kHz and a temporal envelope that was shaped with 1 msec onset and offset ramps. Song pulse rate was varied by changing pulse durations and interpulse intervals in equal amounts to maintain a 50% duty cycle. While pulse rates were varied, total stimulus duration was maintained at a 1 second duration. In total, eight different pulse rate songs were used as test stimuli (20 to 90 pps, in increments of 10 pps). These digital signals were converted to analog signals using the digital-to-analog converter of the National Instruments data acquisition device, amplified using an audio amplifier (Crown XLS1002 Drive Core 2, USA), attenuated with programmable attenuators (Tucker Davis Technologies System 3 PA5, USA), and broadcast through the speakers. Speakers were calibrated to 75 dB SPL (rel. 20 μPa, fast, Z-weighting) at the location of the fly’s head using a probe microphone (Brüel & Kjær, Type 4182, Denmark) connected to a sound level meter (Brüel & Kjær, Type 2250, Denmark).

### General Experimental Protocol

Prior to behavioral experiments, flies were anesthetized on ice for 5 mins and subsequently attached to a tether using a custom low-melting point wax (combination of bee’s wax, rosin, and Sally Hansen wax kit). After this mounting procedure, flies were allowed to acclimate to the testing environment for a period of 15-30 minutes before behavioral testing. Walking phonotaxis experiments were conducted in the dark under IR illumination so that walking responses could be monitored via the digital display on the rear of the high-speed camera equipped with a macro lens (Nikon AF MICRO NIKKOR 105mm f/2.8 D, Japan).

Song pulse rate preferences at different ambient temperatures were tested within subjects. Testing for each subject started at a temperature (21, 25, or 30 °C) that was randomly determined. A 50 pps 1-second control song was presented from the left speaker, followed by the presentation of the same control song from the right speaker. Test songs that varied in pulse rates were initially presented from one speaker for the first 0.5 sec and then switched and presented from the other speaker for the remaining 0.5 sec of stimulus broadcast. Whether the left or right speaker was leading in the stimulus broadcast was chosen at random. Each pulse rate test stimulus was tested three times, resulting in a total of 24 test stimuli presentations at each temperature condition. The order of test stimuli presentations were fully randomized. A particular temperature treatment condition ended with a final round of testing the control stimulus from the left speaker, followed by the right speaker. This testing procedure was replicated at the other two ambient temperatures.

Walking phonotactic responses were considered to be valid if total walking distance exceeded 1 cm. For every three consecutive non-responses to test stimuli, we tested a control stimulus to ensure that the subject was still motivated to respond. If the subject did not respond to this control stimulus, the experiment ended. This data was considered to be incomplete and was excluded from the data analysis. To control for carry-over effects of hearing a previous stimulus, subjects were given 1 minute of rest before testing the next stimulus.

### Data Analysis

Response latencies were determined as the time between stimulus presentation and the first indication of movement (any changes in x or y pixel values). Steering velocities were determined as changes in x pixel values over time. We calculated cumulative total walking distance as:

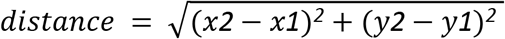

summed across sample points collected during the 1.5 second of data capture. The instantaneous angular heading (in degrees) was determined by converting Cartesian x and y values to polar coordinates by computing the inverse tangent of y divided by x (angular heading = arctan(y/x)). Angular heading was quantified in response to speakers located at ± 45° azimuth (−45°: left speaker, +45°: right speaker). We calculated the mid-response angular heading at the mid-point of stimulus broadcast (0.5 seconds into data capture) which occurred when the broadcast location was switched from the initial location to the subsequent location (e.g. left to right, or right to left). Responses to stimuli where the initial broadcast occurred from the left speaker first were mirrored and reflected across 0° (forward direction). Total walking distance, peak steering velocity, and angular heading were incorporated into calculating the phonotaxis performance index described in [49]. Index values range from 0 to >1. A phonotaxis performance index of 0 indicates poor performance, 1 indicates performance equivalent to responses to the standard song, >1 indicates performance “better” than responses to the control song (e.g., higher steering velocity, longer distance).

We used separate two-way repeated measures ANOVAs to examine the effects of song pulse rate and ambient temperature on 1) response latency, 2) steering velocity, 3) total walking distance, and 4) the phonotaxis performance index. Greenhouse-Geisser corrections were applied where assumptions of sphericity were violated. P-values from multiple post-hoc pairwise comparisons were corrected with Bonferroni adjustments. All statistical analyses were conducted in SPSS Statistics (ver. 19, IBM Corporation, USA). At present, there is no consensus on an inferential statistical approach to analyze repeated measures circular data (but see [57]) for recent developments), thus we opted to use descriptive circular statistics to describe angular heading data in this study.

As PFunc developed by [58] did not allow us to generate appropriately fitted curves to capture preference functions in this current study, we created a MATLAB script that allowed us to generate preference functions based on fitting cubic splines to preferences values. This analysis allowed us to visualize the peak preference and tolerance (or range of pulse rates where the preference is highest) of the preference functions.

## Results

To determine how cricket song pulse rate preferences vary with ambient temperature, we recorded tethered walking phonotaxis from 18 Floridian *Ormia ochracea* using a high-speed spherical treadmill system [56]. Flies were subjected to synthetic crickets that varied in pulse rates (20 to 90 pps, in increments of 10 pps), under three different ambient temperatures (21, 25, and 30°C).

When flies engaged in walking phonotaxis, they responded with mean latencies that ranged from 16.51 msec to 172.53 msec at 21 °C, 15.97 msec to 290.00 msec at 25 °C, and 14.58 msec to 158.33 msec at 30 °C. Response latencies did not vary significantly with pulse rates (rmANOVA: F_7,119_ = 1.685, P = 0.119, η_p_^2^ = 0.09), ambient temperature (F_2,34_ = 0.792, P = 0.461, η_p_^2^ = 0.044), nor was there a significant interaction between pulse rate and ambient temperature (F_14,238_ = 0.953, P = 0.502, η_p_^2^ = 0.053).

We found a significant main effect of pulse rate (rmANOVA: F_7,119_= 11.167, P < 0.001, η_p_^2^= 0.396), temperature (F_2,34_ = 20.275, F < 0.001, η_p_^2^ = 0.544), and the interaction between pulse rate and temperature on total walking distance (F_14,238_ = 2.946, P < 0.001, η_p_^2^ = 0.148). Similar to previously published work [49], flies walked significantly more for a range of pulse rates between 40 to 70 pps compared to higher or lower pulse rates (Fig. 1, Table 1). Flies also had a tendency to walk significantly greater distances at 30 °C than compared to 21 °C (F_1,17_ = 27.822, P < 0.001, η_p_^2^ = 0.621), but walking distances did not significantly differ at 21 °C compared to 25 °C (F_1,17_ = 2.249, P = 0.152, η_p_^2^ = 0.117). A difference contrast (reverse helmert) revealed that flies in an ambient temperature of 30 °C responded with greater walking distances at most pulse rates than compared to responses at 21 °C (Table 2). This temperature by pulse rate interaction was less pronounced when comparing responses at 21 °C versus those at 25 °C. At 21°C, walking distances peaked between 40 to 60 pps, reaching a maximum of 6.77 ± 0.44 cm (mean ± SEM) in response to a 60 pps song. At 25 °C (Fig. 1B), total walking distances across most pulse rates were slightly elevated compared to those observed at 21 °C (Fig. 1A). Walking distance peaked between 50 to 70 pps, with a peak distance of 7.67 ± 0.68 cm in response to 70 pps (Fig. 1B). At 30 °C (Fig. 1C), highest walking distances occurred for a range of pulse rates from 50 to 70 pps, with a maximum of 9.45 ± 0.66 cm (mean ± SEM) in response to the 70 pps song.

**Figure 1.**
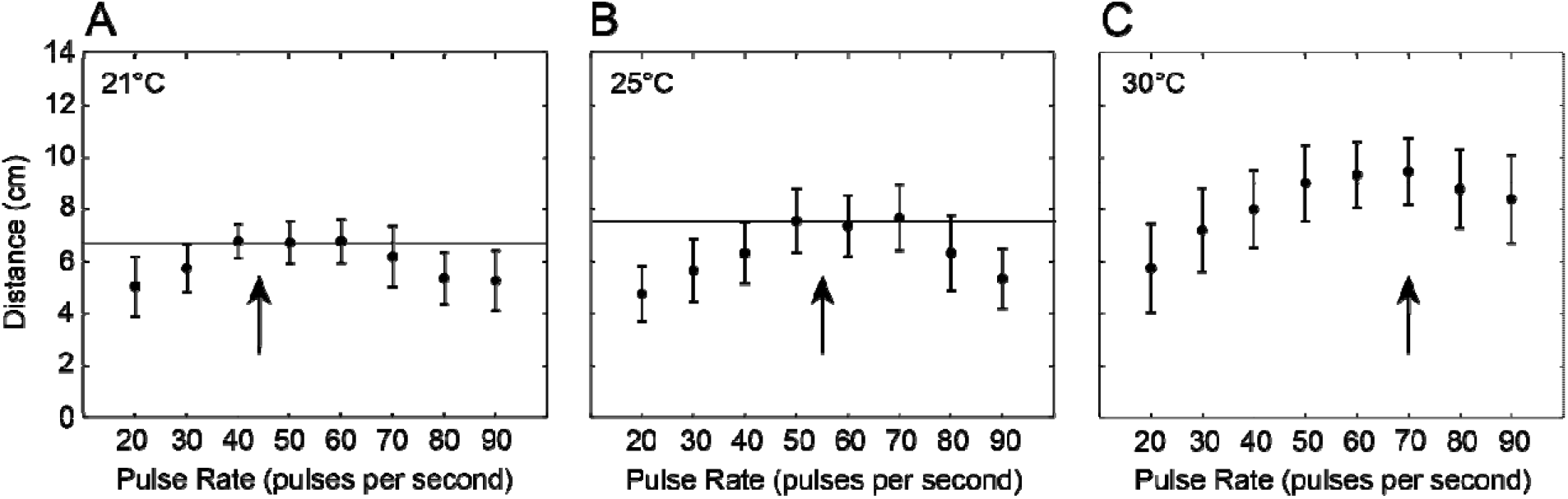
Total walking distance varied with song pulse rate and ambient temperature. Total walking distance (cm) (mean ± SEM) as a function of song pulse rate (pps) at **a)** 21 °C, b) 25 °C, **c)** 30 °C. Arrows denote expected cricket song pulse rate at the specified ambient temperature (21 °C: 44.28 pps, 25 °C: 55.53 pps, 30 °C: 69.34 pps) based on Walker (1962). Horizontal lines indicate total walking distance at 50 pps, which represents the pulse rate of the standard song in the current study.

**Table 1.** Results from analyses of variance of phonotaxis response metrics in response to different pulse rate songs at different ambient temperatures.

| **Response Metric** | **Effect** | **df** | ***F*** | ***P*** | **Partial $\eta^2$** |
| --- | --- | --- | --- | --- | --- |
| Latency | Pulse Rate | 7,119 | 1.685 | 0.119 | 0.090 |
|  | Temperature | 2,34 | 0.792 | 0.461 | 0.044 |
|  | Pulse Rate X Temperature | 14,238 | 0.953 | 0.502 | 0.053 |
| Distance | Pulse Rate | 7,119 | 11.167 | <0.001 | 0.396 |
|  | Temperature | 2,34 | 20.275 | <0.001 | 0.544 |
|  | Pulse Rate X Temperature | 14,238 | 2.946 | <0.001 | 0.148 |
| Steering Velocity | Pulse Rate | 7,119 | 10.224 | <0.001 | 0.362 |
|  | Temperature | 2,34 | 11.46 | <0.001 | 0.389 |
|  | Pulse Rate X Temperature | 14,238 | 3.331 | <0.001 | 0.156 |
| Phonotaxis Index | Pulse Rate | 7,119 | 8.695 | <0.001 | 0.338 |
|  | Temperature | 2,34 | 3.318 | 0.048 | 0.163 |
|  | Pulse Rate X Temperature | 14,238 | 2.247 | 0.07 | 0.117 |

**Table 2.** Post-hoc analysis of difference contrasts to examine temperature effects on response metrics.

| **Response Metric** | **Temperature Comparison** | **Pulse Rate Comparison** | **df** | **F** | **P** | **Partial eta<sup>2</sup>** |
| --- | --- | --- | --- | --- | --- | --- |
| Distance | 25 °C vs 21 °C | 90 vs mean of 80, 70, 60, 50, 40, 30 , 20 | 1,17 | 0.288 | 0.599 | 0.017 |
|  |  | 80 vs mean of 70, 60, 50, 40, 30 , 20 | 1,17 | 1.845 | 0.192 | 0.098 |
|  |  | **70 vs mean of 60, 50, 40, 30, 20** | **1,17** | **4.404** | **0.051** | **0.206** |
|  |  | 60 vs mean of 50, 40, 30 , 20 | 1,17 | 1.532 | 0.233 | 0.083 |
|  |  | **50 vs mean of 40, 30 , 20** | **1,17** | **7.373** | **0.015** | **0.303** |
|  |  | 40 vs mean of 30 , 20 | 1,17 | 0.303 | 0.589 | 0.018 |
|  |  | 30 vs 20 | 1,17 | 0.11 | 0.745 | 0.006 |
|  | 30 °C vs 21 °C | 90 vs mean of 80, 70, 60, 50, 40, 30 , 20 | 1,17 | 1.969 | 0.179 | 0.104 |
|  |  | **80 vs mean of 70, 60, 50, 40, 30, 20** | **1,17** | **4.874** | **0.041** | **0.223** |
|  |  | **70 vs mean of 60, 50, 40, 30, 20** | **1,17** | **8.053** | **0.011** | **0.321** |
|  |  | 60 vs mean of 50, 40, 30 , 20 | 1,17 | 3.571 | 0.076 | 0.174 |
|  |  | **50 vs mean of 40, 30 , 20** | **1,17** | **4.651** | **0.046** | **0.215** |
|  |  | 40 vs mean of 30 , 20 | 1,17 | 1.337 | 0.264 | 0.073 |
|  |  | **30 vs 20** | **1,17** | **5.803** | **0.028** | **0.254** |
| Steering Velocity | 25 °C vs 21 °C | 90 vs mean of 80, 70, 60, 50, 40, 30 , 20 | 1,17 | 0.052 | 0.823 | 0.003 |
|  |  | 80 vs mean of 70, 60, 50, 40, 30 , 20 | 1,17 | 2.306 | 0.146 | 0.114 |
|  |  | 70 vs mean of 60, 50, 40, 30, 20 | 1,17 | 3.266 | 0.087 | 0.154 |
|  |  | **60 vs mean of 50, 40, 30 , 20** | **1,17** | **7.37** | **0.014** | **0.291** |
|  |  | **50 vs mean of 40, 30 , 20** | **1,17** | **4.2** | **0.055** | **0.189** |
|  |  | 40 vs mean of 30 , 20 | 1,17 | 0.121 | 0.732 | 0.007 |
|  |  | 30 vs 20 | 1,17 | 0.049 | 0.828 | 0.003 |
|  | 30 °C vs 21 °C | 90 vs mean of 80, 70, 60, 50, 40, 30 , 20 | 1,17 | 2.632 | 0.122 | 0.17 |
|  |  | **80 vs mean of 70, 60, 50, 40, 30 , 20** | **1,17** | **9.791** | **0.006** | **0.005** |
|  |  | **70 vs mean of 60, 50, 40, 30, 20** | **1,17** | **18.542** | **<0.001** | **0.084** |
|  |  | 60 vs mean of 50, 40, 30 , 20 | 1,17 | 3.243 | 0.089 | 0.153 |
|  |  | 50 vs mean of 40, 30 , 20 | 1,17 | 1.661 | 0.214 | 0.507 |
|  |  | 40 vs mean of 30 , 20 | 1,17 | 0.091 | 0.786 | 0.352 |
|  |  | 30 vs 20 | 1,17 | 3.684 | 0.071 | 0.128 |
| Phonotaxis Index | 25 °C vs 21 °C | 90 vs mean of 80, 70, 60, 50, 40, 30 , 20 | 1,17 | 0.005 | 0.946 | 0 |
|  |  | 80 vs mean of 70, 60, 50, 40, 30 , 20 | 1,17 | 0.812 | 0.38 | 0.046 |
|  |  | **70 vs mean of 60, 50, 40, 30, 20** | **1,17** | **5.979** | **0.026** | **0.26** |
|  |  | 60 vs mean of 50, 40, 30 , 20 | 1,17 | 1.531 | 0.233 | 0.083 |
|  |  | **50 vs mean of 40, 30 , 20** | **1,17** | **11.437** | **0.004** | **0.402** |
|  |  | **40 vs mean of 30 , 20** | **1,17** | **4.243** | **0.055** | **0.2** |
|  |  | 30 vs 20 | 1,17 | 0.246 | 0.626 | 0.014 |
|  | 30 °C vs 21 °C | 90 vs mean of 80, 70, 60, 50, 40, 30 , 20 | 1,17 | 0.51 | 0.485 | 0.029 |
|  |  | 80 vs mean of 70, 60, 50, 40, 30 , 20 | 1,17 | 3.603 | 0.075 | 0.175 |
|  |  | **70 vs mean of 60, 50, 40, 30, 20** | **1,17** | **4.203** | **0.056** | **0.198** |
|  |  | 60 vs mean of 50, 40, 30 , 20 | 1,17 | 2.639 | 0.123 | 0.134 |
|  |  | 50 vs mean of 40, 30 , 20 | 1,17 | 0.493 | 0.492 | 0.028 |
|  |  | 40 vs mean of 30 , 20 | 1,17 | 0.494 | 0.492 | 0.028 |
|  |  | 30 vs 20 | 1,17 | 3.889 | 0.065 | 0.186 |

We also found a significant main effect of pulse rate (rmANOVA: F_1,119_ = 10.224, P<0.001 η_p_^2^=0.362), temperature (F_1,34_ = 11.46, P<0.001, η_p_^2^ =0.362), and the interaction between pulse rate and temperature on average peak steering velocity (F_14,234_=3.331, P <0.001, η_p_^2^=0.156). In general, flies exhibited significantly increased average peak steering velocities for a range of pulse rates between 50 to 70 pps compared to higher or lower pulse rates (Fig. 2, Table 1). Flies exhibited greater peak steering velocities at 30 °C than compared to 21 °C (F_1,17_=13.263, P=0.002, η_p_^2^=0.438), but did not differ for 21 °C compared to 25 °C (F_1,17_=0.034, P =0.855, η_p_^2^=0.002). Analysis of difference contrasts revealed that flies responded with greater peak steering velocities in response to 60 and 50 pps songs relative to all other pulse rate songs at 25 °C compared to responses at 21 °C (Table 2). When steering responses at 30 °C were compared to those at 21 °C, elevated steering responses occurred only at the higher pulse rates of 70 and 80 pps (Table 2). At 21 °C, steering velocity peaked between 40 to 60 pps, reaching a maximum at 50 pps song (5.176 ± 0.29 cm/s). At 25 °C, steering velocity was greatest from 50-70 pps, and reached a peak at 60 pps song (5.444±0.33 cm/s). At 30 °C, peak steering velocity was greatest from 60 to 80 pps, with a peak at the 70 pps song (6.459±0.45 cm/s).

**Figure 2.**
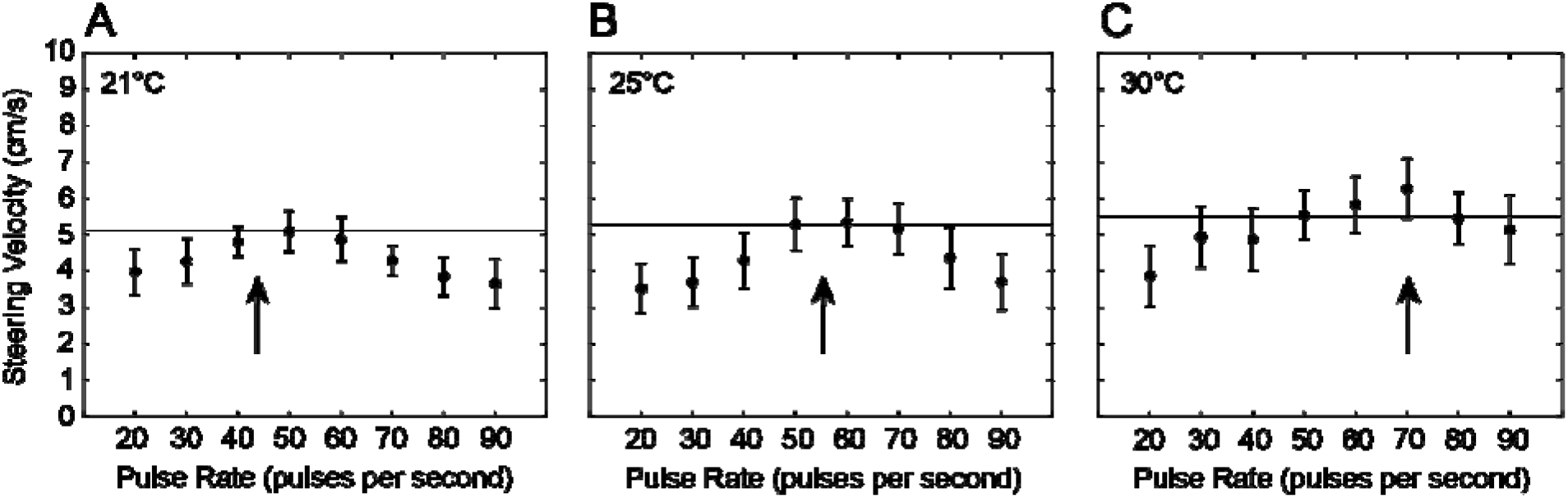
Peak steering velocity varied with song pulse rate and ambient temperature. Peak steering velocity (cm/sec) (mean ± SEM) as a function of song pulse rate (pps) at a) 21 °C, b) 25 °C, c) 30 °C. Arrows denote expected cricket song pulse rate at the specified ambient temperature (21 °C: 44.28 pps, 25 °C: 55.53 pps, 30 °C: 69.34 pps) based on Walker (1962). Horizontal lines indicate peak steering velocity at 50 pps, which represents the pulse rate of the standard song in the current study.

Mean angular heading appeared to vary with song pulse rate preferences (Fig. 3). There was a greater tendency for flies to perform walking phonotaxis with angular headings that approached 45° (location of the speaker) at the most preferred pulse rates (40-70 pps). For example, mean angular heading at 21 °C for pulse rates at 40, 50, and 60 pps were 31.60°, 95% CI [51.09°, 75.33°], 31.71°, 95% CI [50.05°, 76.78°], and 29.69°, 95% CI [41.48°,77.26°], respectively. At 25 °C, mean angular headings for the peak pulse rates, 50 and 60 pps were 32.19°, 95% CI [ 47.26°, 81.51°] and 30.97°, 95% CI [44.89°,79.01°]. At 30 °C, the mean angular heading at 60 pps was 31.45°, 95% CI [ 51.26°, 74.53°] and 30.00°, 95% CI [47.34°, 72.64°] at 70 pps. However, angular headings were highly variable in response to most pulse rates and ambient temperature levels, suggesting that the accuracy in localizing the sound source may not depend on temperature.

**Figure 3.**
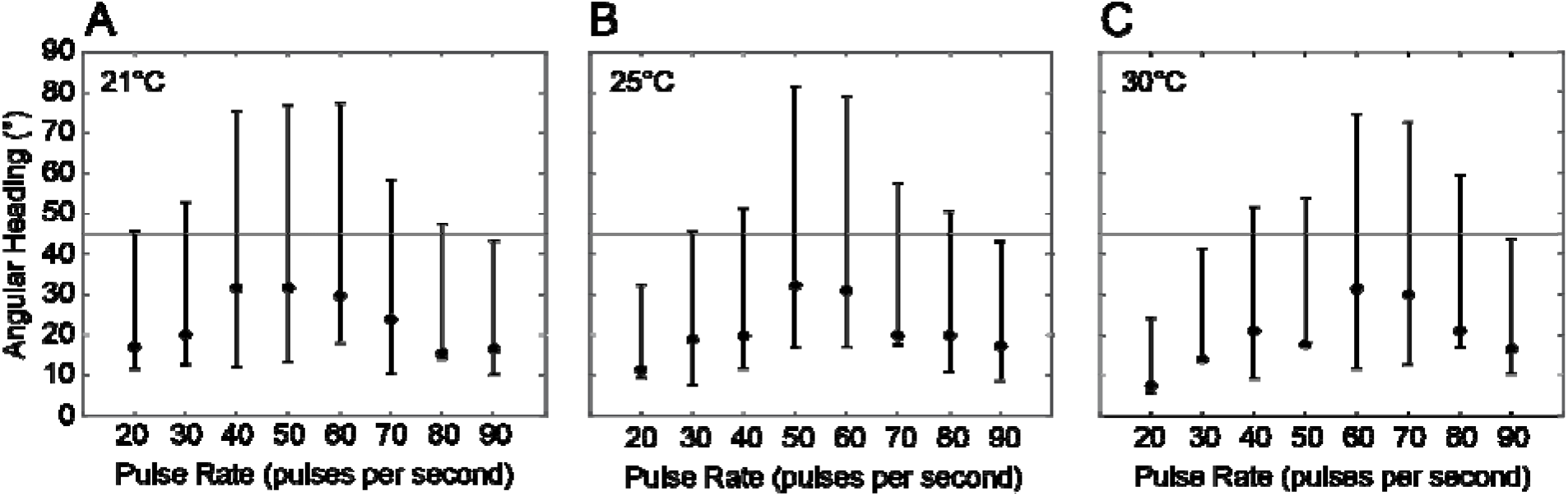
Angular heading varied with song pulse rate and ambient temperature. Angular heading (mean ± 95% confidence interval) as a function of song pulse rate (pps) at **a)** 21 °C, b) 25 °C, **c)** 30 °C. Horizontal lines indicate speaker location at 45° azimuth.

We examined the effects of temperature on pulse rate preferences and overall phonotactic performance using our previously described phonotaxis performance index [49]. There was a significant main effect of pulse rate (F_7,119_=8.695, P <0.001, η_p_^2^=0.338), temperature (F_2,34_=3.318, P =0.048, η_p_^2^=0.163), and the interaction between pulse rate and temperature on the phontaxis performance index (F_14,238_=2.247, P <0.07, η_p_^2^=0.117). Flies had significantly higher phonotaxis performance index scores for a range of pulse rates between 40 to 80 pps compared to those at higher or lower pulse rates (Fig. 4, Table 1). Analysis of difference contrasts revealed that when 25 °C was compared to 21 °C, flies responded with higher phonotactic performance at most intermediate pulse rate values (e.g. 70, 50, 40 pps, see Table 2). At 30 °C compared to 21 °C, peak performance was highest in response to 70 pps relative to all lower pulse rates (Table 2).

**Figure 4.**
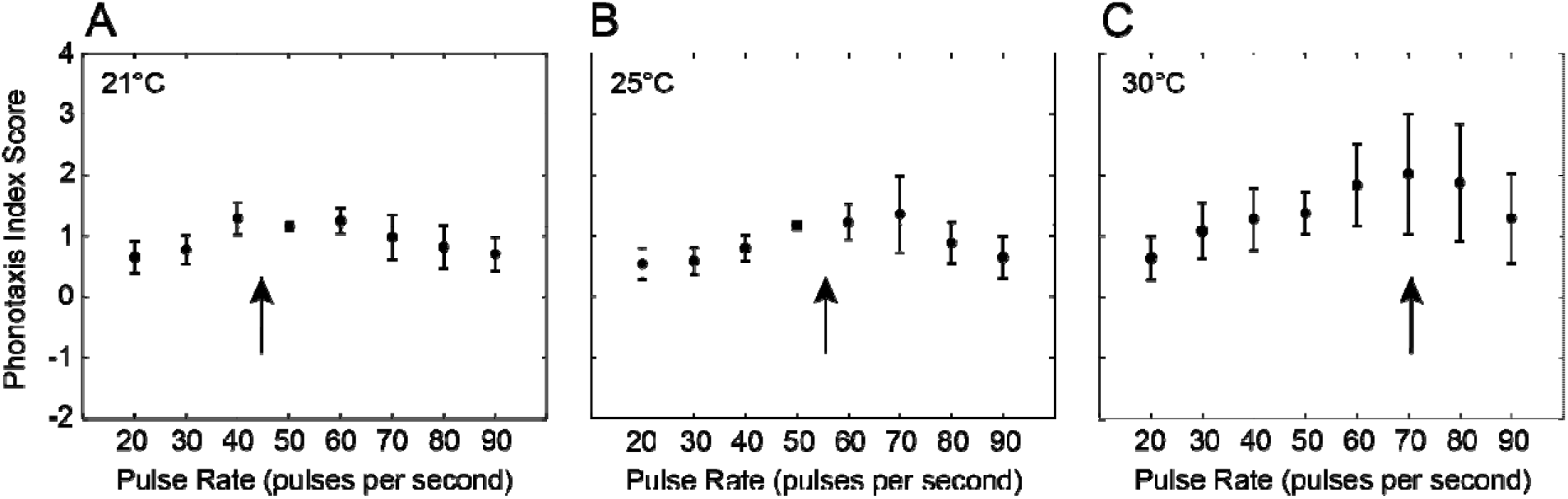
Phonotaxis performance index score varied with song pulse rate but not ambient temperature. Phonotaxis performance index scores (mean ± SEM) as a function of song pulse rate (pps) at **a)** 21 °C, b) 25 °C, **c)** 30 °C. Arrows denote expected cricket song pulse rate at the specified ambient temperature (21 °C: 44.28 ppa, 25 °C: 55.53 ppa, 30 °C: 69.34 pps) based on Walker (1962). Horizontal lines indicate phonotaxis performance index score at 50 pps, which represents the pulse rate of the standard song in the current study.

To characterize function valued signal preferences that relate to finding suitable host crickets, we developed a custom MATLAB script with similar functionality to PFunc [58]. This MATLAB script allowed us to fit cubic splines to phonotaxis performance index scores as a function of song pulse rates (Fig. 5). The fitted preference functions across individuals varied in shape. For example, some flies had preference functions that were linear with peak phonotaxis performance at extreme pulse rate values (e.g. 20 or 90 pps). Some individuals had bimodal preference functions, and others had preference functions that had peaks at intermediate values. Of the 18 flies, 61% (11/18) of flies demonstrated a shift in the peak pulse rate preference to higher pulse rate songs at higher ambient temperatures, 39% (7/18) of flies demonstrated an increase in tolerance as seen by an increase in the broadness of their pulse rate preference functions.

**Figure 5.**
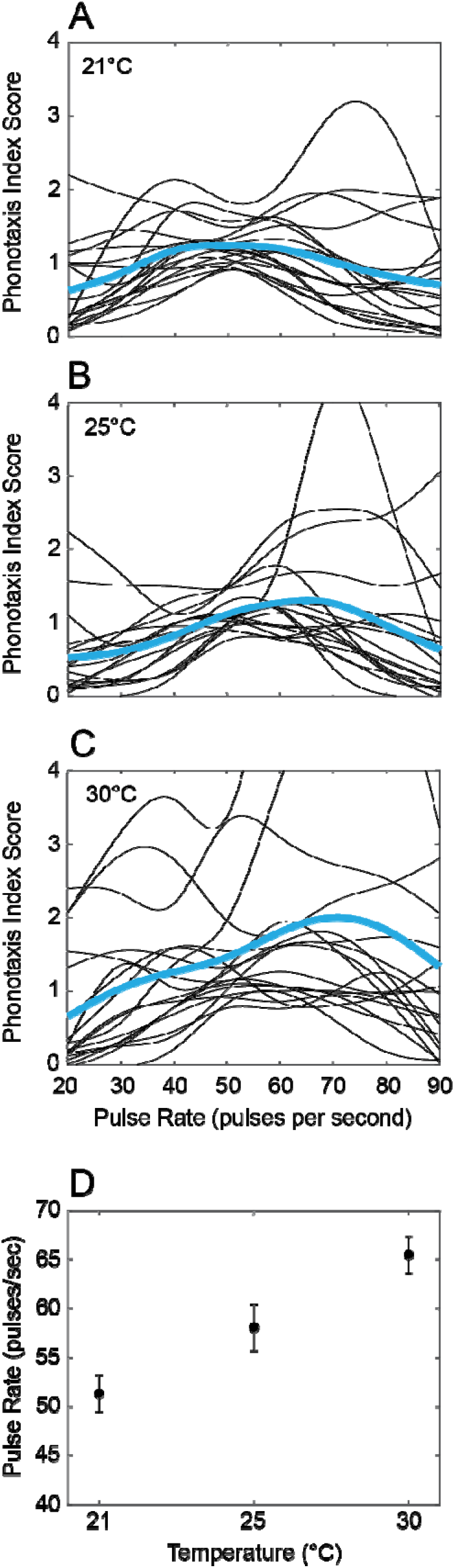
Pulse rate preference functions based on phonotaxis performance index scores as a function of ambient temperature. Preference functions were developed as cubic splines fitted to phonotaxis performance index scores as they varied with song pulse rates at **a)** 21 **°C, b)** 25 **°C, c)** 30 **°C.** Black lines represent individual preference functions (n=18 flies) while blue lines represent mean preference pulse rate function. Axes in panels **a), b),** and **c)** scaled to emphasize mean preference functions, which resulted in the cropping of some individual preference curves. Peak pulse rate preference values were extracted from individual curves and the mean±SEM peak pulse rate preference is plotted as a function of ambient temperature in **d).**

Among 72% of the flies tested (13 of 18 flies), peaks were successfully identified from fitted preference functions across pulse rates and all ambient temperature conditions. There was a significant main effect of temperature on the peak in pulse rate preference (rmANOVA: F_2,34_=13.12, P <0.0001, η_p_^2^=0.522). At an ambient temperature of 21 °C, the pulse rate preference function had a peak at 51.31 ± 1.87 pps (mean ± SEM) (Fig. 5A,D). At 25 °C, the peak preference value increased to a significantly higher pulse rate of 58.08 ± 2.36 pps (Fig. 5B,D, F_1,12_=5.047, P =0.044, η_p_^2^=0.296). When the ambient temperature was increased to 30 °C, the peak of the pulse rate preference function increased to 65.46 ± 1.87 pps, which was significantly different from the peak preference exhibited at 21 °C (Fig. 5A,D, F_1,12_=31.97, P <0.001, η_p_^2^=0.727).

Shifts in the peak of the pulse rate preference function as a function of ambient temperature also occurred along with a non-significant, but trending tendency in the broadening of preference functions (F_2,28_=3.008, P =0.066, η_p_^2^=0.177). At 21 °C, tolerance values were 27.86 ± 1.49 (mean ± SEM). These tolerance values increased to 33.92 ± 3.81 and 39.24 ± 3.81 when the ambient temperature was increased to 25 °C and 30 °C respectively.

## Discussion

As temporal features of cricket calling songs vary with ambient temperature [20,21,25,59], we sought to determine if temperature-dependent changes in song patterns would affect the ability of *Ormia ochracea* to recognize or alter preferences for the calling songs of suitable host cricket species. This was assessed by examining the effects of ambient temperature on several key walking phonotactic response metrics. With the exception of response latencies, all other metrics (total distance, peak steering velocity, angular heading, phonotaxis performance index) varied with song pulse rate, which is consistent with our previous work [49]. At 21 °C, the magnitude of these response metrics had a tendency to peak at around 50-70 pps, and declined at higher and lower pulse rates. At the highest ambient temperature tested (30 °C), these response metrics generally increased and reached a plateau in response to higher pulse rates, resulting in preference functions with a “high-pass” appearance. Over this same range of ambient temperatures, the calling song pulse rate of *Gryllus rubens* has been demonstrated to increase from 40 to 70 pps [59]. Our results revealed a change in the peak of pulse rate preference functions that suggests a level of “temperature coupling” as seen in intraspecific acoustic communication systems among some orthoperans and anurans [19,22,24,25,36].

Walker [47] performed a series of sound trap experiments in the field to determine if the “acoustic template” of *O. ochracea* varied with ambient temperature. In these experiments, a *Gryllus rubens* calling song with a temperature-dependent pulse rate consistent with the ambient temperature was broadcast from one sound trap, while a calling song with a pulse rate that was temperature adjusted to be either 4 °C above or below (equivalent to 11 pps higher or lower than the temperature appropriate song pulse rate) the ambient temperature was broadcast from the alternative sound trap. Under these experimental conditions, the sound trap broadcasting a pulse rate song consistent with the ambient temperature attracted the most flies, which suggested that the internal acoustic template for song recognition in *O. ochracea* is temperature-dependent. Our current results extend these past findings in two important ways. First, Walker (1993) only tested a limited range of pulse rates (11 pps above and below the preferred pulse rate) while our current study examined a greater range of pulse rate selectivity (20-90 pps) that allows for a function-valued approach in developing signal preference functions. These signal preference functions across different ambient temperatures are consistent with perceptual neurosensory mechanisms involved in song recognition that may vary with ambient temperature. Second, beyond field capture rates, our results are the first to demonstrate how walking phonotaxis response metrics change with temperature.

An increase in ambient temperature caused a shift in the peak of the pulse rate preference functions, along with a slight tendency for an increase in tolerance for a wider range of pulse rates. Consistent with previous findings [49], our current results demonstrate that *O. ochracea* prefers cricket songs with a pulse rate of 50 pps at 21 °C. At higher temperatures, *O. ochracea* walked faster and farther, which contributed to higher phontaxis performance index values in response to generally less preferred higher pulse rate songs (>70 pps). Such increases in these walking response metrics indicate an increase in the overall phonotactic responsiveness and the broadening of acceptable pulse rates that can elicit phonotactic behavior. Together, these temperature-dependent changes resulted in preference functions that were “closed-ended” at 21 °C, but more “open-ended” at 30 °C.

Temperature coupling between signalers and intended receivers may depend on common genetics and neural architecture [36] among conspecifics to adaptively maintain signal recognition between conspecifics across a wide range of ambient temperatures. However, in some instances, temperature coupling seems to be an emergent property of cellular processes that change with temperature in ways that may not contribute to signal recognition [60]. For example, in the short horned grasshopper *Omocestus viridulus*, increases in ambient temperature causes chirp duration preferences to diverge from actual song durations produced at higher temperatures [61]. In the acoustic moth *Achroia grisella*, the pulse-pair rate produced by males increases with temperature (18-30 °C tested), and female pulse-pair rate response threshold also increases with temperature, but the male pulse-pair rates were all above the recognition threshold at all temperatures tested [60]. In contrast, gravid female *O. ochracea* are not genetically coupled with their host crickets. Any temperature dependent changes in song recognition and signal preferences may be based on common temperature effects on underlying cellular processes, which may include auditory neurons that are involved in song pattern recognition [62]. Such mechanisms underlying the receiver’s psychology (including unintended receivers), would be under strong selection to either be tuned, or have sufficient boardness. As reproductive success in *Ormia ochracea* is highly dependent on recognizing and localizing suitable host cricket calling songs, *O. ochracea’s* acoustic template for host cricket song recognition should be broad and flexible to account for small changes in temporal features across different ambient temperatures.

Several gaps in our understanding of song pattern recognition, and signal preferences in *O. ochracea* still remain. It is known that *O. ochracea* occurs across parts of Mexico, in the southern US, and on some Hawaiian Islands [63]. In each of these regions, *O. ochracea* are behaviorally specialized to prefer specific host cricket species [48] and it is clear that some populations of *O. ochracea* may recognize and utilize multiple host cricket species [47,64,65]. Species-specific differences in cricket songs are largely based on the temporal patterning of sound pulses [63]. How fine-scale temporal features contribute to song preferences and song recognition is unknown. For example, Floridian *O. ochracea* prefers cricket songs within a range of pulse rates (45-50 pps) [47,49]. This pulse rate preference may be based on a preference for a particular pulse duration, interval between pulses, or different combinations of durations and intervals that make up a particular pulse period [66]. Whether different populations of *O. ochracea* evaluate the same temporal features (e.g. durations, intervals, pulse periods) but prefer a different range of values, and how temperature may affect song recognition across populations, remains to be determined. In the lab, we have demonstrated that the “acoustic template” of Floridan *O. ochracea* may be broad enough to allow for host song recognition across a range of temperatures, but future work should address to what extent *O. ochracea* encounter this range of temperature during their phonotactic approach in nature.

## Acknowledgements

We thank Eric S. Cole for feedback on an earlier version of this manuscript; past and present members of the Lee Lab of Neural Systems and Behavior for animal care; and Wade W. Schulz and Kurtis W. Johnson for technical support. N.L. thanks Mi Jung Kim for support throughout this project. N.L. also dedicates this manuscript to his mother Tung Moy Lee and father Chee Wing Lee, both Canadian immigrants from China, that worked hard to provide N.L. with an education that made this work possible. This work was supported by a U.S. National Science Foundation CAREER grant to N. Lee. (IOS-2144831), the St. Olaf College Collaborative Undergraduate Research and Inquiry (CURI) Program, and the St. Olaf TRIO McNair Scholars Program.

## Author Contributions

Conceptualization, K.J.J., N.L.; Methodology, N.L.; Software, N.L.; Investigation, K.J.J., I.A., J.G., E.S., L.J.A., N.L.; Resources, N.L.; Data Curation, J.A.D., N.L; Writing - Original Draft, K.J.J., J.A.D., N.L.; Writing -Review and & Editing, J.A.D., N.L.; Visualization - K.J.J., J.A.D, N.L.; Funding Acquisition, N.L.

